# AFMnanoSALQ: An Accurate Detection Framework for Semi-Automatic Labeling and Quantitative Analysis of α-Hemolysin Nanopores Using Intensity-Height Cues in HS-AFM Data

**DOI:** 10.1101/2025.02.26.640237

**Authors:** Thuyen Vinh Tran Nguyen, Ngoc Quoc Ly, Ngan Thi Phuong Le, Hoang Duc Nguyen, Kien Xuan Ngo

## Abstract

High-Speed Atomic Force Microscopy (HS-AFM) enables imaging of biological structures and dynamics with nanometer spatial and millisecond temporal resolution. AFM images contain three-dimensional (3D) surface information, comprising two-dimensional (2D) lateral (x-y) and one-dimensional (1D) height (z) encoded in pixel intensity. This dynamic structure poses significant challenges for instance boundary detection and morphological analysis. To address this, we develop AFMnanoSALQ, a feature-driven computational framework for semi-automatic labeling and quantitative (SALQ) detection and morphological measurement of HS-AFM data. Unlike conventional methods that rely solely on either visual or geometric features for 2D boundary detection, AFMnanoSALQ integrates both to extract 3D morphology. It requires neither annotated data nor intensive training, enabling fast deployment at minimal cost. With performance comparable to typical deep-learning models, AFMnanoSALQ facilitates semi-automatic labeling, making it a practical tool for preliminary data inspection and accelerating the creation of training datasets. As a case study, we focus on α-hemolysin (αHL), a β-barrel pore-forming toxin secreted by *Staphylococcus aureus*, using both synthetic and experimental AFM data. AFMnanoSALQ provides a foundation for future deep learning studies, enabling both dataset generation and cross-validation between feature-driven and data-driven approaches.

## 1 Introduction

HS-AFM[1] provides nanometer (nm) resolution (∼1 nm laterally; ∼0.1 nm vertically) with millisecond (ms) temporal imaging. The resulting AFM images capture 3D surface information, where spatial position (x, y) is defined by lateral coordinates and height (z) is represented by pixel intensity. However, complex noise and heterogeneity in AFM images across different conditions make analysis challenging, requiring tailored methods that effectively exploit both spatial and height features.

Recent advances have transformed HS-AFM surface images into 3D atomic conformations using rigid or flexible fitting techniques [2-10], while AlphaFold 2 and 3 have revolutionized structure prediction from single conformations to complex assemblies involving proteins, nucleic acids, ligands, ions, and modified residues[11-14]. However, most methods often assume isolated targets, whereas real HS-AFM images typically contain multiple or heterogeneous conformations. Simulation-based approaches that generate pseudo-AFM images from Protein Data Bank (PDB)^1^ have been explored, but without incorporating force-field constraints, they risk inaccurate 3D fitting due to missing physical interactions[15][16]. This highlights the need for robust data preprocessing and accurate instance boundary detection.

Although deep learning achieves state-of-the-art performance, it relies heavily on large annotated datasets and requires finetuning across imaging conditions. To address some of those limitations, we propose AFMnanoSALQ, a feature-driven computational framework for instance boundary detection and morphological measurement of HS-AFM data. Unlike conventional methods that rely solely on visual or geometric features for 2D boundary detection, AFMnanoSALQ integrates both to extract 3D morphology. It requires neither annotated data nor intensive training, allowing deployment at minimal cost. AFMnanoSALQ facilitates semi-automatic labeling, making it a practical tool for preliminary data inspection and for accelerating the creation of training datasets.

As a case study, we focus on α-hemolysin (αHL)[17-20], a pore-forming toxin that disrupts host cell membranes. Despite extensive study, no effective vaccine exists, underscoring the need for in-depth analysis. HS-AFM images of αHL exhibit distinct geometric features—mushroom-shaped and pore structures—well suited for feature-driven approaches. These characteristics support instance boundary detection and morphological measurement, facilitating the analysis of pore formation stages.

The framework comprises three stages. Preprocessing stage removes noise and imaging artifacts[21] using Non-local Mean[22] and DeStripe[23] filters, with parameter tuning to preserve structural features. Instance Boundary Detection stage targets αHL “kernels,” whose geometry varies across developmental stages, using a three-step protocol: (i) Kernel Detection uses the Hessian matrix to identify candidate regions; (ii) Kernel Verification leverages the αHL geometry and the orthogonal, top-down HS-AFM imaging setup to perform multi-level clustering based on height, confirming candidates consistently surrounded by circular or arc-shaped boundaries; (iii) Curvature Boundary Extraction refines pixel-level edges for precise boundary delineation. Finally, in the quantification stage, morphological parameters (e.g., perimeter, area, volume) are computed via derivative and integration-based, providing a comprehensive quantitative description of αHL’s conformations.

As with most feature-driven approaches, the effectiveness of AFMnanoSALQ depends on the structural characteristics of the target. Customizing feature extraction may be necessary when adapting to different molecular systems.

## 2 Related Work

This section reviews traditional and deep learning methods for denoising and instance boundary detection in AFM images, highlighting their strengths and limitations.

### 2.1 Denoising

AFM generates images through the interaction between the cantilever probe and the sample surface, making it prone to point noise, stripe artifacts, and scanning distortions[21]. Point noise appears as isolated specks, stripe noise as linear artifacts, and distortions often arise from scanning issues. To mitigate these, both traditional and deep learning denoising methods have been widely explored.

Traditional computational methods provide diverse strategies for denoising while preserving structural details. *Morphology-based*[24] techniques (e.g., erosion, dilation) remove isolated noise pixels but risk losing fine details. The *Median filter*[24] reduces impulse noise by replacing each pixel with the median of its neighborhood, maintaining edge sharpness. *Non-Local Means (NLM)*[22] averages similar pixels across the image to lower noise while retaining critical features. *Gaussian filters*[24] are fast but may smooth out details. *DeStripe*[23] targets stripe noise in the frequency domain with minimal impact on image content. *Wavelet-based denoising*[25] removes noise via soft thresholding in the frequency space, preserving sharp transitions. *Low-Rank Recovery*[26] models the image as a sum of low-rank (clean) and sparse (noise) component, effectively isolating structured noise like stripes, though at higher computational cost.

**Table 1.**
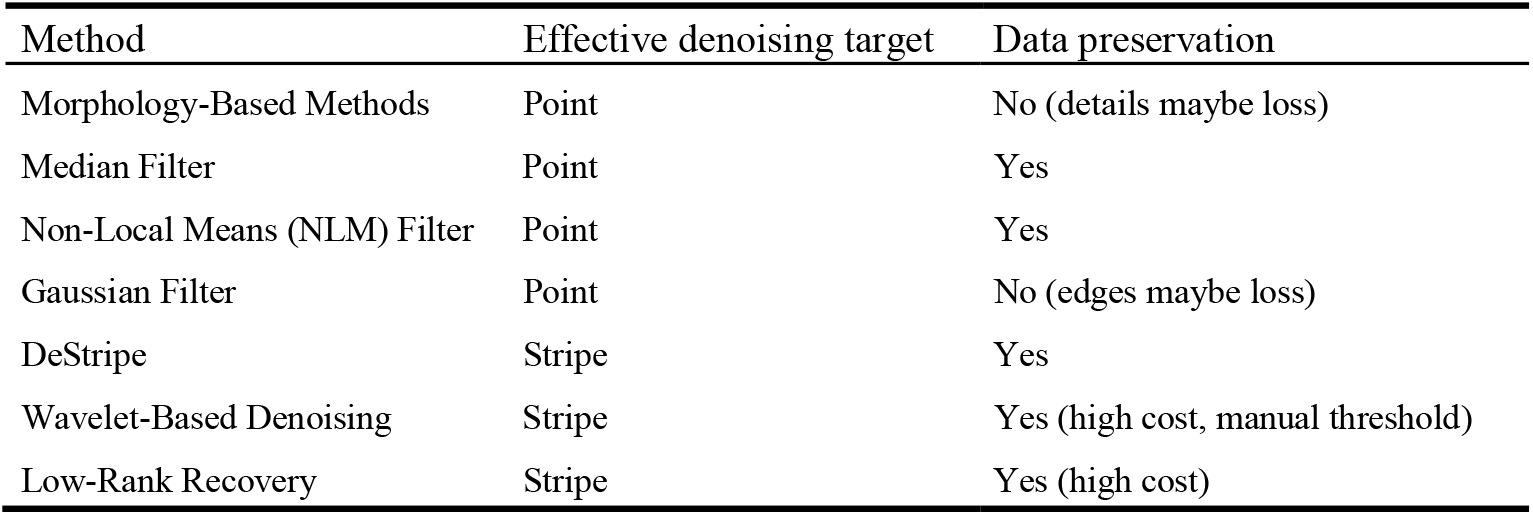
Evaluation and comparison between *traditional computational methods*.

| Method | Effective denoising target | Data preservation |
| --- | --- | --- |
| Morphology-Based Methods | Point | No (details maybe loss) |
| Median Filter | Point | Yes |
| Non-Local Means (NLM) Filter | Point | Yes |
| Gaussian Filter | Point | No (edges maybe loss) |
| DeStripe | Stripe | Yes |
| Wavelet-Based Denoising | Stripe | Yes (high cost, manual threshold) |
| Low-Rank Recovery | Stripe | Yes (high cost) |

Deep learning methods have also been applied to HS-AFM denoising. *MPRNet*[27] uses multi-stage residual learning and multi-scale fusion to remove complex noise while preserving fine details. *HINet*[27] extracts hierarchical multi-scale feature for robust denoising. *Uformer*[27] uses a U-shaped transformer with self-attention to capture long-range dependencies and address both point and stripe noise. *Restormer*[27], optimized with multi-head self-attention for reduced complexity, excels at eliminating noise while maintaining structural integrity. According to [27], *Restormer* is the most effective, while *HINet* is the most efficient for denoising AFM images.

Traditional methods are efficient, interpretable, and require no labeled data, allowing real-time processing with controllable noise–detail trade-offs. However, they may over-smooth and obscure nanoscale features under complex noise patterns. Deep learning approaches can effectively remove complex noise patterns but require large annotated datasets and computational resources. This study focuses on exploring the potential of traditional methods. In future work, deep learning techniques will be experimentally evaluated and compared with the traditional approaches for further enhancement.

### 2.2 Instance Boundary Detection

This study categorizes related methods for accurate object delineation in HS-AFM images along two axes: traditional vs. deep learning methods, and visual (intensity-based) vs. geometric (3D topology–based) features.

**Table 2.** Categorization table of *instance boundary detection methods*.

|  | Visual (Intensity-based) | Geometric (3D Topology-based) |
| --- | --- | --- |
| Traditional Methods | - Edge-Based<br>- Blob Detection | - Watershed (Ridge/Valley Detection)<br>- Geodesic Active Contours & Level Set |
| Deep Learning Methods | - SegNet<br>- UNet<br>- YOLOv8-Seg | - Depth-aware CNNs<br>(include visual intensity) |

Traditional methods rely on explicit mathematical operations and heuristics:

- *Edge-Based[24]*: detect intensity gradients using operators like Sobel or Canny. While computationally simple and interpretable, they are highly sensitive to noise and low contrast in HS-AFM images, causing fragmented or incomplete boundaries.
- *Blob Detection[28]*: uses the Hessian matrix or Laplacian of Gaussian to identify blob-like regions representing object centers. However, variability in object shape and intensity necessitates careful parameter tuning.
- *Watershed[24]*: the best ridge/valley segmentation[29], treats the image as a topographic surface, flooding from local minimum to define boundaries. Though effective for contiguous regions, it is prone to over-segmentation in noisy images unless marker-controlled variants are applied with careful parameter tuning.
- *Geodesic Active Contours with Level Set[30]*: segments objects by iteratively evolving contours along topological features, considering height levels. Effective for complex shapes but sensitive to initialization and computationally intensive. Given enough annotated data, deep learning achieves superior performance:
- *SegNet[31]*: uses an encoder–decoder architecture to capture spatial hierarchies via convolutional layers, enabling pixel-wise segmentation even in noisy environments.
- *Unet[32]*: Combines low-level spatial details with high-level semantics via skip connections, enabling accurate boundary delineation—well-suited for HS-AFM images where fine structural details are critical.
- *YOLOv8-Seg[33]*: Offers fast instance-level segmentation but may lack boundary precision at nanoscale and relies heavily on labeled data.
- *Depth-Aware CNNs[34]*: use height or depth maps as additional input channels for topology-aware segmentation, commonly applied to RGB-D images.

In comparison, traditional methods are efficient and interpretable, while deep learning offers high accuracy but requires large labeled datasets, high computational cost and lacks transparency. However, most existing approaches tend to rely on either visual intensity or geometric cues alone. *Given that HS-AFM encodes both, our method integrates both visual intensity and geometric height cues feature types to fully exploit the data’s structure, enabling semi-automatic labeling and quantitative analyses*. Our approach accelerates dataset creation for deep learning models, particularly in analyzing αHL.

## 3 Method

AFMnanoSALQ performs denoising, instance boundary detection, and measurement of morphological parameters (**Scheme 1**) to analyze the conformational dynamics of αHL, which forms in a lipid membrane containing cholesterol, using HS-AFM imaging.

**Scheme 1:.**
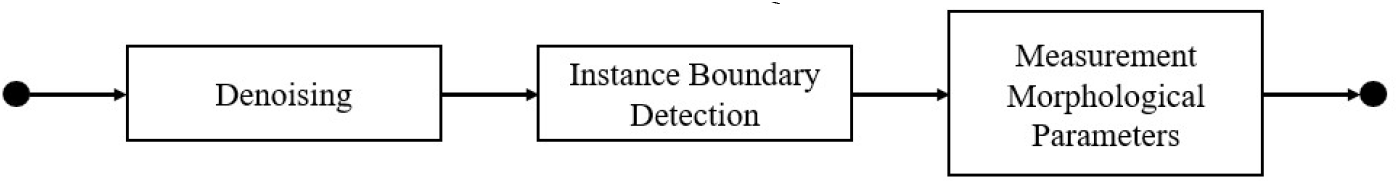
A workframe of AFMnanoSALQ.

### Dataset

This study uses the dataset from [35][36]. The αHL[17-19], a pore-forming toxin secreted by *Staphylococcus aureus*, disrupts host cell membranes. Despite extensive research, no vaccine exists, underscoring the need for better diagnostics[20]. Understanding the oligomerization and pore formation process is essential for vaccination.

The dataset (**Fig. 1, Appendix 1, 2**) is organized into three distinct categories:

**Fig. 1.**
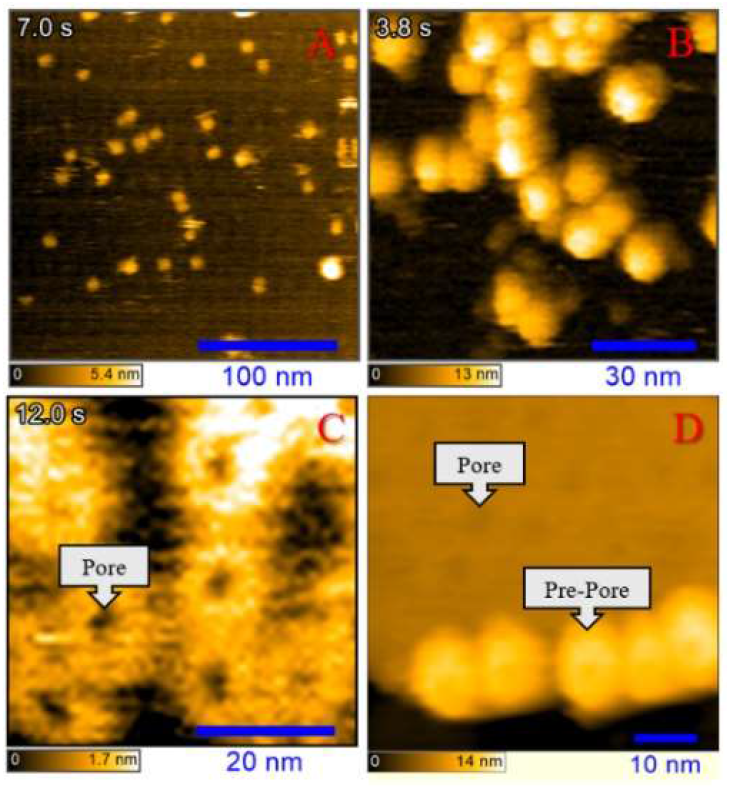
Representative HS-AFM images illustrating developmental stages of αHL. (**A**) ***Intermediate Structures***; (**B**) ***Pre-Pore Structures***; (**C**) ***Pore Structures***; (**D**) ***Pre-Pore and Pore Structures***. The top-left displays the elapsed time (in seconds) from the start of the imaging sequence, enabling temporal correlation of dynamic events. The bottom-left shows the vertical (height-z) color-scale bar, which converts the color-coded height data into nanometers for precise vertical measurements. The bottomright presents the lateral (length and width) scale bar, translating pixel dimensions into nanometers to accurately quantify lateral dimensions.

1. **Intermediate Structures**: This includes monomers, dimers, trimers, tetramers, pentamers, hexamers … formation during the oligomerization process (**Fig. 1A**).
2. **Pre-Pore Structures**: Characterized by mushroom-like heptamer structures, formed on membrane surface with a height of approximately 10 nm (**Fig. 1B, D**).
3. **Pore Structures**: the transmembrane domain of prepore conformation inserted into the lipid membrane, causing the height to decrease from 10 nm to around 6 nm in AFM images (**Fig. 1C, D)**.

All AFM image series in the dataset are stored in ASD format and analyzed by the UMEX Viewer software, developed by Kenichi Umeda (Kanazawa University). To standardize the processing workflow, HSAFM images are exported from ASD to PNG format with embedded metadata, ensuring accurate real-world measurements.

### Problem Definition

The key challenge is accurately detecting target “kernels”— oligomers, mushroom-shaped pre-pores[18][19], and pores—from HS-AFM images, followed by reliable morphological quantification. For each αHL developmental stage, the objectives are:

- *Intermediate Structures*: detect oligomers (kernel) to measure perimeter, area, height…; enabling volume estimation during oligomerization process.
- *Pre-Pore Structures*: extract the characteristic mushroom-shaped region (kernel) and compute its height to evaluate the progression of membrane penetration.
- *Pore Structures*: isolate post-membrane penetration pores (kernel), quantify perimeter, area, volume… to elucidate the structural transition from pre-pore to pore states.

Mathematically, given a sequence of HS-AFM images capturing each stage in the development of αHL, the goal is to accurately detect target “kernel” and compute their morphological parameters in physical units (nanometers). Formally, let

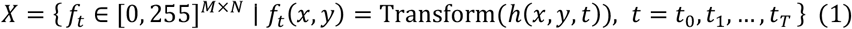

where *X* is the set of HS-AFM images observed over time; *M* and *N* denote the image dimensions (length and width); *f*_t_ is an image frame acquired at a specific time *t*; and *f*_t_(*x, y*) = Transform(*h*(*x, y, t*)) denotes the pixel value, where *h*(*x, y, t*) represents the physical surface height at coordinates (*x, y*) in the image captured at time *t*, and Transform(⋅) maps these height values into grayscale intensity.

The objective is to detect boundary of each predefined “kernel” in each stage. Let

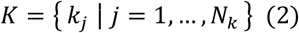

be the set of segmented targets obtained from an image *x* ∈ *X*. A preprocessing operator *P*_*pre*_ is first applied to enhance the image (e.g., smoothing, grayscale converting, denoising), followed by a segmentation operator *S*:

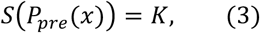

Subsequently, for each segmented kernel *k* ∈ *K*, we define a measurement function

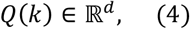

which computes morphological parameters in physical units (nanometers).

In summary, AFMnanoSALQ preprocesses HS-AFM images via *P*_*pre*_, detects the target “kernels” via *S* and computes morphological parameters via *Q* to quantitatively analyze “kernels” of various αHL throughout its developmental stages (**Fig. 2**).

**Fig. 2.**
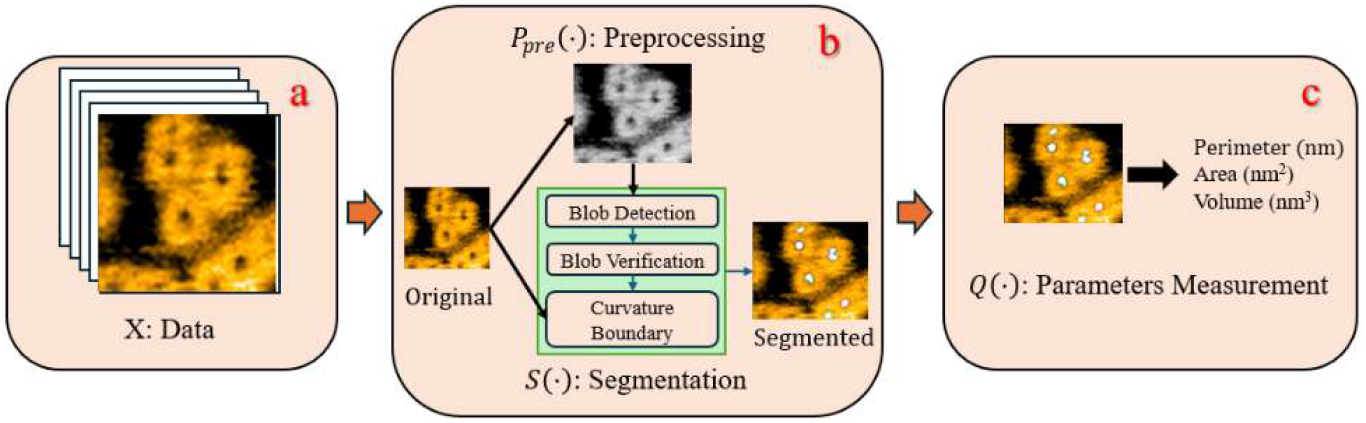
Overview of AFMnanoSALQ for analyzing HS-AFM images of αHL. **(a)** Dataset *X* comprises HS-AFM images encoding height via color, capturing αHL dynamics over time. **(b)** Function *P*_*pre*_(⋅) performs preprocessing (e.g., smoothing, grayscale converting, denoising) and function *S*(⋅) detects instance boundary of “kernel” in conformations (e.g., oligomers, mushroom-shaped pre-pores, pores). **(c)** Function *Q*(⋅) computes morphological parameters (e.g., perimeter, area, volume, height, depth) in physical units (nm). Together, these components form a comprehensive pipeline for quantitative analysis of αHL developmental stages in HS-AFM data.

#### 3.1 Denoising

A two-pronged approach was adopted to address HS-AFM primary noises: Non-local Mean[22] for point noise and DeStripe[23] for stripe noise. To avoid detail loss or artifacts, denoised images are used only for instance boundary detection; measurements are taken from original images using the detected boundary masks for accurate quantification. In this study, traditional methods are prioritized for their efficiency, interpretability, and independence from labeled data—aligning with the goal of semi-automatic labeling, thereby to establish a foundation for developing deep-learning in the future.

#### 3.2 Instance Boundary Detection

In image processing, a *blob* is defined as a localized region with consistent intensity or topology that stands out from its surroundings, often exhibiting near-circular, elliptical, or more generally, strongly curved boundaries. In HS-AFM images, αHL structures appear differently through various developmental stages —dome-like oligomers, mushroom-shaped pre-pores, and cylindrical pores—exhibit precisely blob characteristics, making Hessian-based blob detection suitable, as it detects regions with strong curvature via second-order derivatives.

### Kernel Detection

Candidate kernel centers are identified by detecting blob-like regions via Hessian matrix computation at each pixel (*x, y*) of the image *I* (**Fig. 3d**):

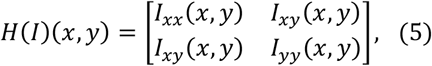

where *I*_xx_ and *I*_yy_ are the second-order derivatives along the *x* and *y* directions, and *I*_xy_ is the mixed derivative.

**Fig. 3.**
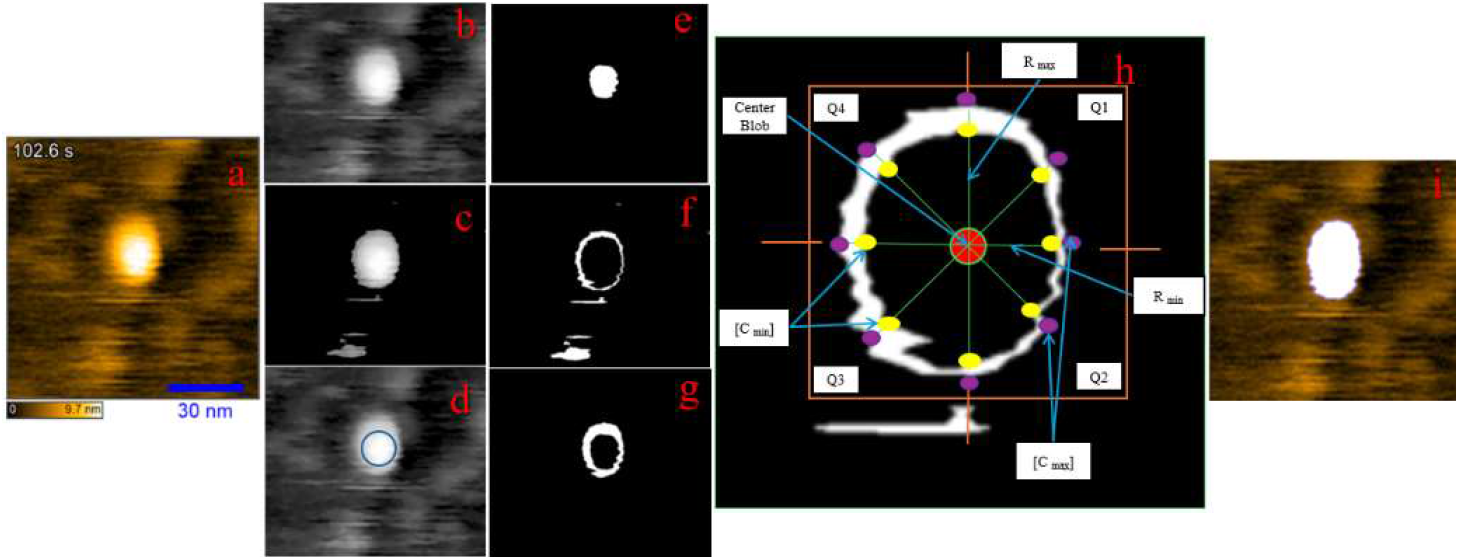
Segmentation workflow for αHL in the pre-pore stage. **(a)** Original image; **(b)** Grayscale and cropped image (scalebar and elapsed time removed); **(c)** Denoised image with background removed; **(d)** Blob detection identifying potential blobs; **(e**,**f**,**g)** Multi-level set clustering based on height; **(h)** Blob verification, “Center Blob” marks potential blob center, [C_min_] and [C_max_] denote the initial and extended raycast points used to determine the minimum (R_min_) and maximum (R_max_) radius, and Q1, Q2, Q3, Q4 represent quadrants for assessing spatial distribution; **(i)** Curvature boundary extraction precisely delineates and highlights the object boundary.

The Hessian matrix, which involves second-order derivatives, is well-suited for HS-AFM images, which exhibit smooth gradient variations. Unlike first-order derivatives (e.g., Canny, Sobel) that often miss subtle boundaries, second-order derivatives enhance curvature sensitivity, improving structure detection in this context.

The Hessian determinant det(*H*(*I*)(*x, y*)) is calculated and identifies local maxima as candidate blobs. These regions, characterized by high determinant values, indicate isotropic intensity variations consistent with blob-like structures.

### Kernel Verification

Following initial detection, candidate blobs are verified by leveraging αHL morphology and top-down AFM imaging, which encodes height via color channels. Multi-level set partitioning is performed via K-means clustering across height levels (**Fig. 3e, f, g**).

For each height level, a valid candidate blob is expected to be surrounded by a nearly circular shape. In such a shape, the enclosing pixels satisfy the circle equation

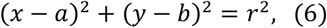

where (*a, b*) represents the candidate blob’s center. Considering that blobs have a finite size, the center coordinates are allowed to vary within specified bounds, i.e., *x* ∈ [*x*_min_, *x*_max_] and *y* ∈ [*y*_min_, *y*_max_], and the radius *r* vary within [*r*_min_, *r*_max_](**Fig. 3h**).

To determine *r*_min_ and *r*_max_, raycasting is performed from blob center along eight principal directions. The first intersections with the non-background form set *C*_min_. Rays are extended until re-hit the background, and intersections form set *C*_max_. Distances from blob center to all points in *C*_min_ and *C*_max_ are computed, and rays deviating more than 1.5 times the median length of the set (either shorter or longer) are discarded. Finally, *r*_min_ and *r*_max_ are defined as the minimum and maximum distances between blob center to the remaining points in *C*_min_ and *C*_max_, respectively (**Fig. 3h**).

Conceptually, using filtered *C*_min_, region growing is applied to aggregate adjacent pixels satisfying the circle equation, yielding set *X*. The area around the blob center is divided into four quadrants, and pixels in *X* are assigned accordingly. A balanced distribution across quadrants is checked to avoid directional bias despite geometric fit. Then, connected component analysis is applied to assess the density and coherence of *X*. A candidate is accepted as a valid kernel if *X* forms a compact, near-circular region across multiple height levels. This multi-step verification ensures only well-defined kernels are retained for further analysis (**Fig. 3h**).

Operationally, to enhance efficiency, region growing is avoided in favor of localized evaluation using a sliding mask from each point in *C*_min_ and *C*_max_. Boundary is verified via non-background pixel ratio in a local mask. The mask can be adaptively shifted based on heuristics. Additionally, parallel processing is employed to evaluate multiple candidate blobs across height levels simultaneously.

### Curvature Boundary Extraction

After confirming a blob, its boundary is delineated using the Watershed algorithm, which is well-suited for HS-AFM data as it encodes 3D structures and can be treated as a topographic map, enabling a landscape-based interpretation, where valleys indicate regions and ridges boundaries (**Fig. 3i**).

Watershed partitions a gradient image *G* into catchment basins based on local minima. With domain Ω and local minima {*m*_i_}, each catchment basin *C*_i_ is defined as:

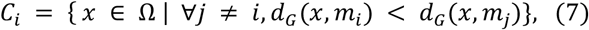

where *d*_G_(*x, m*) is a distance metric that incorporates the gradient values along paths from *x* to *m*. The watershed lines *W* are pixels on the boundaries between basins:

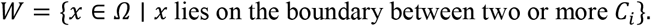

Although Watershed is prone to over-segmentation by noises, constraining it to verified blob regions minimizes spurious boundaries, enhances delineation accuracy.

In summary, curvature boundary extraction combines a modified watershed algorithm—constrained to verified blob regions. This multi-stage approach ensures accurate and robust boundary delineation for reliable morphological measurements. Given the 3D nature of HS-AFM and the cylindrical shape of αHL pores, tuning the Watershed flooding level parameters allows separation of base, shaft, and top of pore—highlighting the algorithm’s suitability for αHL analysis.

### 3.3 Measurement

Once the “kernels” have been accurately detected, key morphological parameters like the perimeter and the area of each kernel are computed.

### Perimeter Calculation

Let 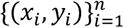 denote the sequential boundary points. The perimeter *P* is then computed using a discrete approximation by summing the Euclidean distances between successive points:

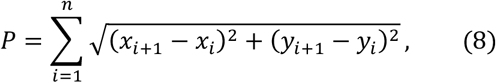

with the convention that (*x*_n+1_, *y*_n+1_) = (*x*_1_, *y*_1_) to ensure the boundary is closed.

### Area Calculation

The area *A* of the object is determined via integration over the region *D* enclosed by the boundary. In the continuous domain, this is expressed as:

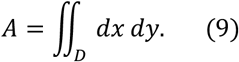

The area can be computed from the boundary points via Green’s theorem [37]:

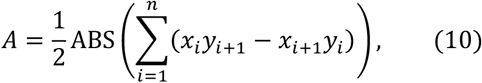

### Conversion to Physical Unit

After computing perimeter and area in pixel units, conversion to nanometers is required for physical interpretation. This involves detecting the embedded scale bar (**Fig. 1**) based on its position and geometry, followed by extracting text indicating the physical length using Google Tesseract OCR.

Let *D*_pixel_ be the scale bar length (in pixels) and *D*_physical_ be its physical length (in nanometers) read by OCR. The conversion factor *s* is computed as:

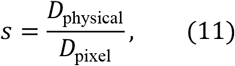

Using this factor, the perimeter *P*_pixel_ and area *A*_pixel_ are converted by:

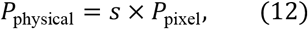

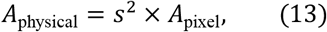

This approach ensures morphological measurements are accurately translated into nanometer-scale dimensions, essential for quantitative HS-AFM analysis.

### Volume Calculation

After boundary detection, volume is calculated using the height information encoded in the HS-AFM image. For each pixel (*x, y*) within the segmented region *D*, the volume *V* is the integral of the height difference between the local height *h*(*x, y*) and baseline height *h*_0_, typically taken at the boundary.

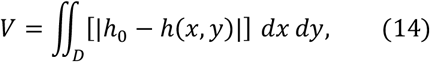

In digital images, this integral is approximated by a discrete sum:

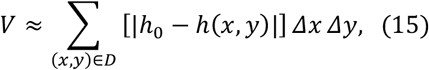

where Δ*x* and Δ*y* are the physical dimensions (in nanometers) of each pixel.

Due to pixel resolution limits, raw height data *h*(*x, y*) may exhibit abrupt transitions that do not accurately represent the true continuous surface. To address this, the height map is refined through smoothing. The local height gradient is estimated using finite difference methods to approximate the derivative ∇*h*(*x, y*). Then, a smoothing filter (e.g., Gaussian) is applied to generate a refined height map 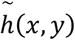 that more accurately captures continuous surface variations.

The volume is then recalculated using the smoothed height values:

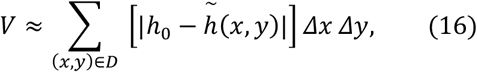

After computing the volume in pixel units, the color-to-nanometer conversion factor is extracted by reading the color scalebar in image (**Fig. 1**). This factor is combined with the previously obtained pixel-to-nanometer conversion factor, and the volume is converted using the formula:

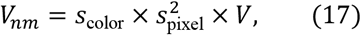

Where 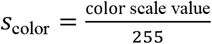 and *s*_pixel_ is the pixel-to-nanometer conversion factor.

## 4 Experiments

### 4.1 Dataset Description

The dataset comprises HS-AFM image sequences categorized as follows:

- Intermediate Structures: 25 sequences, totaling 6,500 images.
- Pre-Pore Structures: 20 sequences, totaling 4,000 images.
- Pore Structures: 50 sequences, totaling 10,000 images.

### 4.2 Denoising Evaluation

As outlined in the Methods, this study prioritizes traditional denoising approaches to align with the goal of semi-automatic labeling without reliance on annotated data.

Peak Signal-to-Noise Ratio (PSNR) is the primary metric for assessing denoising performance by measuring the mean squared error (MSE) between the denoised image and the original noisy image. The PSNR is defined mathematically as:

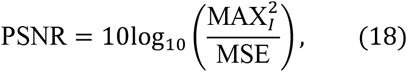

where MAX_*I*_ represents the maximum pixel value. The MSE is calculated as:

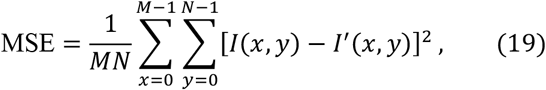

with *I*(*x, y*) and *I*^’^(*x, y*) denoting the pixel value at position (*x, y*) in the original image and the denoised image, respectively; *M* and *N* being the image dimensions.

PSNR yields results in decibel (dB) units, quantifies denoising effectiveness— higher PSNR value indicates that the processed image contains less noise.

Table 3 shows that Non-local Mean (NLM) achieves the highest and most stable PSNR. DeStripe (and NLM + DeStripe) yield lower PSNRs on original images but perform relatively better after cropping out the scale bar and elapsed text. Accordingly, NLM is adopted as the primary denoising method, while using DeStripe (recommended in [38]) or its combination with NLM selectively for pre-cropped images.

**Table 3.**
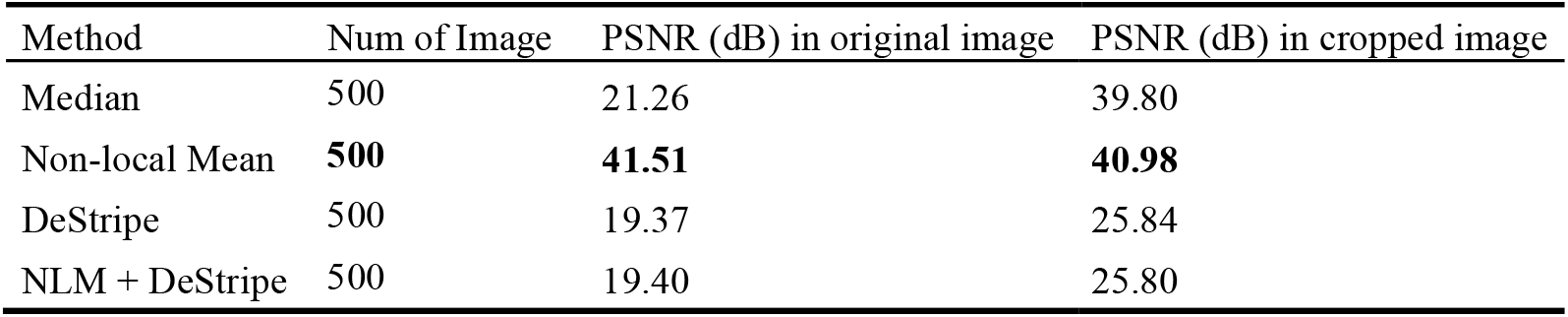
PSNR evaluation for denoising methods on original images (include scalebar/text) and cropped images (cropped 8% top, 10% bottom to remove scalebar/text), as shown in **Fig.1**.

### 4.3 Instance Boundary Detection Evaluation

The semi-automatic labeling capability and standalone applicability of AFMnanoSALQ for preliminary data inspection are systematically evaluated in this section. To establish benchmarks, UNet and YOLOv8-Seg were selected as baseline models due to their popularity and broad applicability. UNet is well-established for its accuracy in pixel-wise segmentation and boundary delineation, while YOLOv8-Seg offers state-of-the-art performance in real-time object detection with integrated mask prediction. This enables benchmarking AFMnanoSALQ against established approaches and facilitates indirect comparison with more advanced models through standard reference points [39-41]. The evaluation is conducted in three main categories:

#### Labeling Evaluation

Two labeling strategies were evaluated for accuracy:

- **SA (Semi-Auto):** Labels were generated using AFMnanoSALQ, which integrates both 2D-intensity and 3D-morphology-height features from HS-AFM images. To further enhance accuracy, labels were manually refined using AFMnano3DR, a visualization tool that treats AFM images as 3D point clouds for surface re-construction (**Appendix 1**). Refinement focused on recovering missed pores caused by ambiguity or partial truncation, particularly at image edges.
- **M (Manual)**: Manual annotations were performed using QuPath [42], relying exclusively on 2D-intensity information from HS-AFM images. Structural or height-based cues were not incorporated.

For individual αHL conformations at various stages, a training subset was selected discretely from image sequences across the dataset to ensure diversity and avoid near-duplicate frames, then independently labeled using two strategies, forming **Label_SA** and **Label_M**. Three annotators independently labeled the dataset, with disagreements resolved by majority vote; Cohen’s kappa for inter-annotator agreement reached 0.89. Labeling consistency was evaluated using three criteria: Object Count (total detected objects), Matched Objects (number of matching objects), Matched IoU (IoU of matched pairs) and Annotation Time (average time to annotate a single image). **Table 4** shows over 90% consistency between strategies and 56% annotation time reduction with **SA**.

**Table 4.** Comparison of Label_SA and Label_M averaged over subsets from individual á HL conformations (intermediate, pre-pore, pore)

| Labeled Set | Object Count | Matched Object | Matched IoU | Annotation Time |
| --- | --- | --- | --- | --- |
| Label_SA | 7.42 | 7.06 | 92% | 3.6 minutes |
| Label_M | 7.37 |  |  | 8.2 minutes |

#### Object Detection Performance

Detection accuracy was evaluated using precision, recall, and F1-score. An independent set of 100 carefully annotated images, labeled by three domain experts (Cohen’s κ = 0.94), was used as groundtruth (**Valid_GT**).

UNet and YOLOv8-Seg were independently trained using Label_SA and Label_M, split at the sequence level into training (80%) and testing (20%) sets, ensuring all frames of a sequence resided entirely in one set to avoid data leakage. To mitigate limited data, k-fold cross-validation was applied across sequences. The splits shared identical image compositions; only the annotation strategies differed. This yielded 4 models: UNet_SA, UNet_M, YOLOv8-Seg_SA, and YOLOv8-Seg_M, with performance evaluated by **Valid_GT**, which also benchmarked the standalone accuracy of AFMnanoSALQ.

As shown in **Table 5**, YOLOv8-Seg_SA achieved the highest F1-score, while UNet_SA yielded the highest IoU. Models trained with Train_SA achieved higher recall and competitive precision. This can be attributed to the SA method’s integration of second-order derivatives, which is highly sensitive to changes in intensity and curvature—combined with height information, enhancing object detection capability. In contrast, manually annotated based on 2D-intensity, focused on clearly visible objects, resulting in fewer labeling errors but also omitting potential objects. Notably, AFMnanoSALQ, when used independently, provided comparable detection performance, underscoring its utility as an efficient semi-automatic annotation as well as practical tool for preliminary data inspection that significantly reduces manual labeling effort.

**Table 5.**
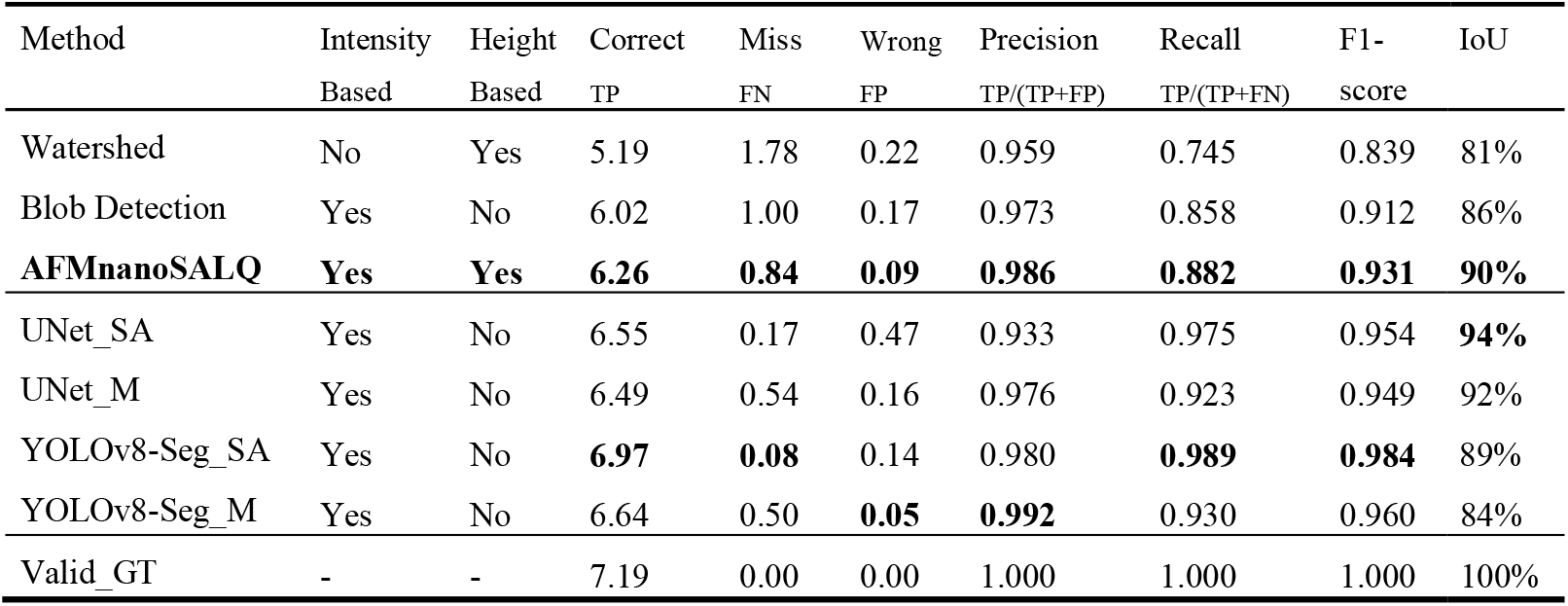
Object Detection Evaluation averaged over subsets from individual αHL conformations (intermediate, pre-pore, pore)

| Method | Intensity<br>Based | Height<br>Based | Correct<br>TP | Miss<br>FN | Wrong<br>FP | Precision<br>TP/(TP+FP) | Recall<br>TP/(TP+FN) | F1-score | IoU |
| --- | --- | --- | --- | --- | --- | --- | --- | --- | --- |
| Watershed | No | Yes | 5.19 | 1.78 | 0.22 | 0.959 | 0.745 | 0.839 | 81% |
| Blob Detection | Yes | No | 6.02 | 1.00 | 0.17 | 0.973 | 0.858 | 0.912 | 86% |
| **AFMnanoSALQ** | **Yes** | **Yes** | **6.26** | **0.84** | **0.09** | **0.986** | **0.882** | **0.931** | **90%** |
| UNet_SA | Yes | No | 6.55 | 0.17 | 0.47 | 0.933 | 0.975 | 0.954 | **94%** |
| UNet_M | Yes | No | 6.49 | 0.54 | 0.16 | 0.976 | 0.923 | 0.949 | 92% |
| YOLOv8-Seg_SA | Yes | No | **6.97** | **0.08** | 0.14 | 0.980 | **0.989** | **0.984** | 89% |
| YOLOv8-Seg_M | Yes | No | 6.64 | 0.50 | **0.05** | **0.992** | 0.930 | 0.960 | 84% |
| Valid_GT | - | - | 7.19 | 0.00 | 0.00 | 1.000 | 1.000 | 1.000 | 100% |

#### Object Measurement Accuracy

detection quality is assessed by shape and quantitative measurements, with shape accuracy measured via Intersection over Union (IoU) between groundtruth annotations and detected regions. (**Table 5**).

Average morphological parameters obtained from AFMnanoSALQ are compared with reference values from PDB and prior studies on αHL to validate detection accuracy, measurement precision, and consistency with established literature (**Table 6**).

**Table 6.** Pore Measurement Evaluation.

| Method | Num of Sample | Average Perimeter (nm) | Average Area (nm<sup>2</sup>) |
| --- | --- | --- | --- |
| Manual by UMEX Viewer | 1109 | 11.58 ± 1.14 | 9.31 ± 1.82 |
| **AFMnanoSALQ** | 1686 | 12.68 ± 2.11 | 10.56 ± 3.58 |
| PDB,[17],[18],[19] | - | 9.425 ± 5.03 | 9.08 ± 7.54 |

Notably, AFMnanoSALQ measured αHL height at ∼9.94 nm in the pre-pore stage and ∼6.46 nm in the pore stage (measured from the membrane upward), closely matching published data[19][35]. Direct volume computation on pore-stage images is hindered by tip-sample dilation, resolution limits, and scale conversion errors. To address this, the average pore area and height value across stages were used to indirectly estimate the internal volume, yielding ∼70 nm^3^, consistent with prior studies [17],[18],[19].

**Appendix 2** shows AFMnanoSALQ outputs as qualitative evidence of effectiveness.

## 5 Conclusion

AFMnanoSALQ facilitates semi-automatic labeling and quantification with rapid and reasonably accurate detection, supporting preliminary data inspection and significantly accelerating training dataset creation for deep-learning models. As a framework tailored for analyzing αHL conformations, it performs robust instance boundary detection and quantitative analysis on HS-AFM images by extracting both visual intensity and geometric height cues beyond the currently available methods, effectively addressing challenges such as noise and heterogeneous boundary and pore structures. Experimental results on αHL confirm its effectiveness and adaptability. However, as with most feature-driven approaches, performance depends on the structural characteristics of the target and parameter tuning is occasionally required to achieve optimal results; adapting to different molecules may require customizing feature extraction. This study provides a foundation for future work, with labeled outputs from AFMnanoSALQ serving to support deep learning models and enabling large-scale investigation on 3D surface or atomic structures through SimHS-AFMfit and dynamics of proteins obtained from HS-AFM data [43].

## Acknowledgments

This research is supported by research funding from Faculty of Information Technology, University of Science, Viet Nam National University – Ho Chi Minh City.

We also gratefully acknowledge the Bio-SPMs Collaborative Research Program at WPI-NanoLSI, Kanazawa University, for providing NTPL and HDN support in imaging and analysis. The authors also acknowledge the financial support from KAKENHI (Japan Society for the Promotion of Science) for KXN (19K06581, 23K05713, 23H02452-01).

## Appendix 1. AFMnano3DR Visualization Tool

In this study, the AFMnano3DR tool is employed to visualize the structure of αHL (**Fig. 4**) directly from HS-AFM images and to reconstruct its 3D surface representation (**Fig. 5**). Beyond visualization, AFMnano3DR is further utilized to refine masks from AFMnanoSALQ semi-automatic labeling and quantification, enhancing the accuracy and reliability of the final annotations.

**Fig. 4.**
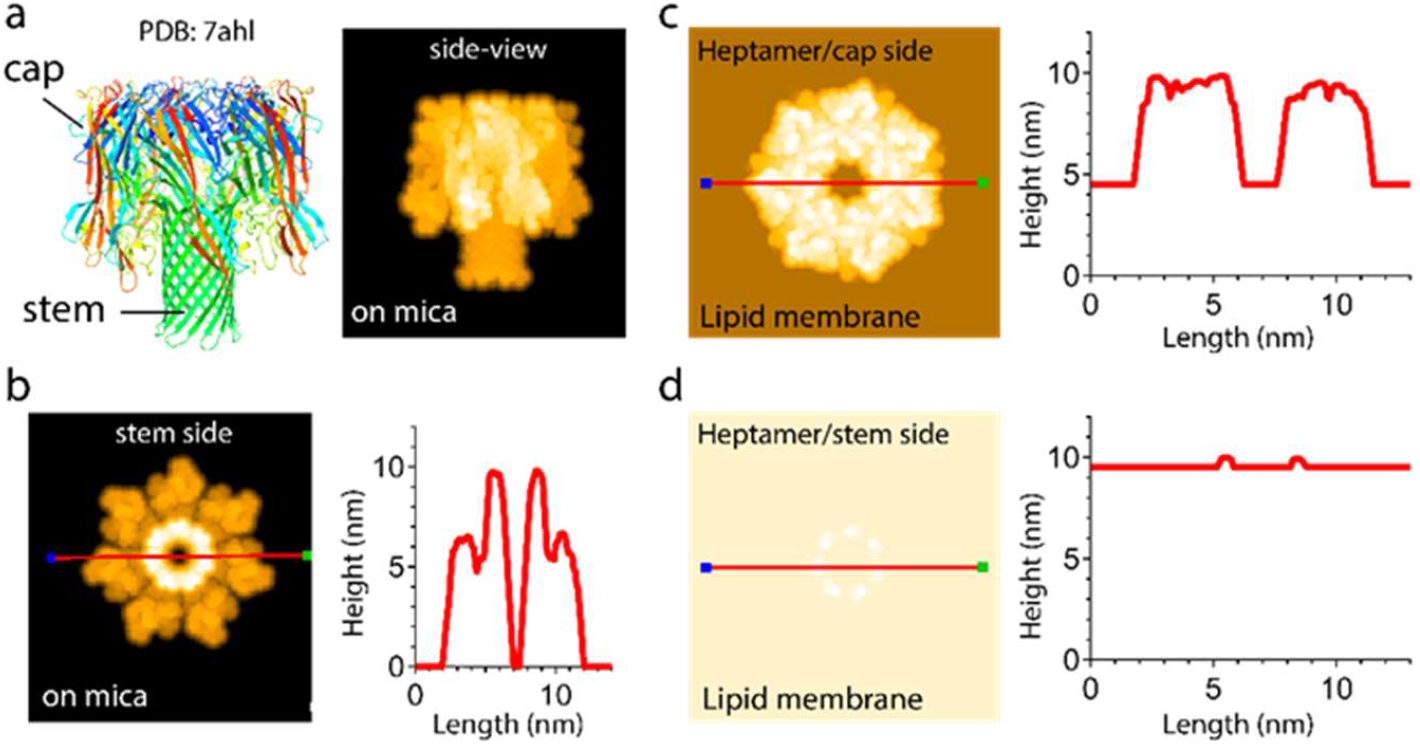
Structural features of the αHL heptameric pore. (a) Domain composition, ribbon model, and pseudo-AFM image of the heptarmeric pore (PDB: 7ahl), (b) Pseudo-AFM image viewed from the stem side (left) with corresponding cross-sectional profile (right), (c, d) Pseudo-AFM images of the pore inserted into the lipid membranes, viewed from cap side and stem side, respectively, along with their cross-sectional profiles.

**Fig. 5.**
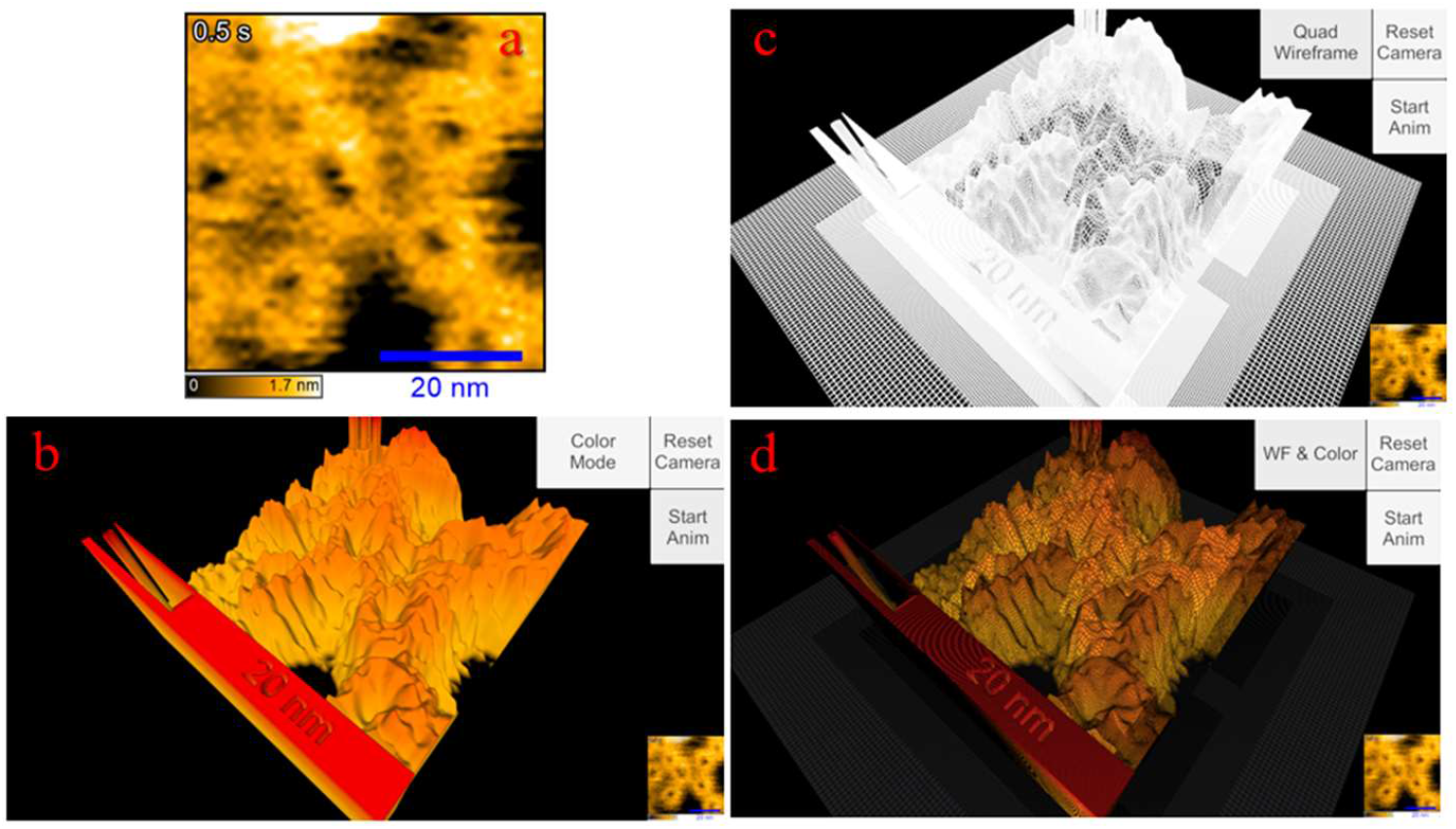
3D surface reconstruction of αHL using AFMnano3DR. **(a)** HS-AFM input image; **(b)** 3D wireframe mesh reconstruction; **(c)** 3D surface mesh reconstruction; **(d)** Combined wireframe and surface view.

AFMnano3DR is a software tool developed to reconstruct 3D surfaces from AFM images. It treats AFM data as a topographic point cloud, where each pixel encodes spatial (x, y) and height (z) information and reconstructs a continuous 3D surface byinferring point connectivity via a custom-designed algorithm. The tool supports generating both triangle-based and quad-based wireframe meshes as well as full surface renderings (**Fig. 5**), enabling structural visualization of nanoscale objects.

The tool includes a wide range of features to support preprocessing and interactive analysis (**Fig. 6, Fig. 7**) such as denoising, filters, morphological operations, region selection, and surface marking using a 3D brush. Users can also generate 3D and 2D masks based on manually marked or algorithmically selected regions.

**Fig. 6.**
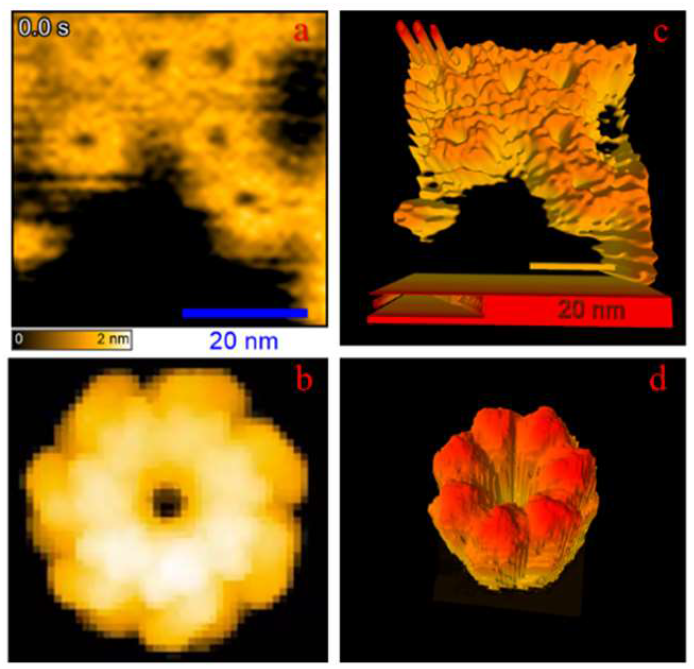
Examples of 3D surface reconstruction using AFMnano3DR. **(a), (b)** Input HS-AFM images; **(c), (d)** Corresponding 3D surface meshes reconstructed from **(a)** and **(b)**.

**Fig. 7.**
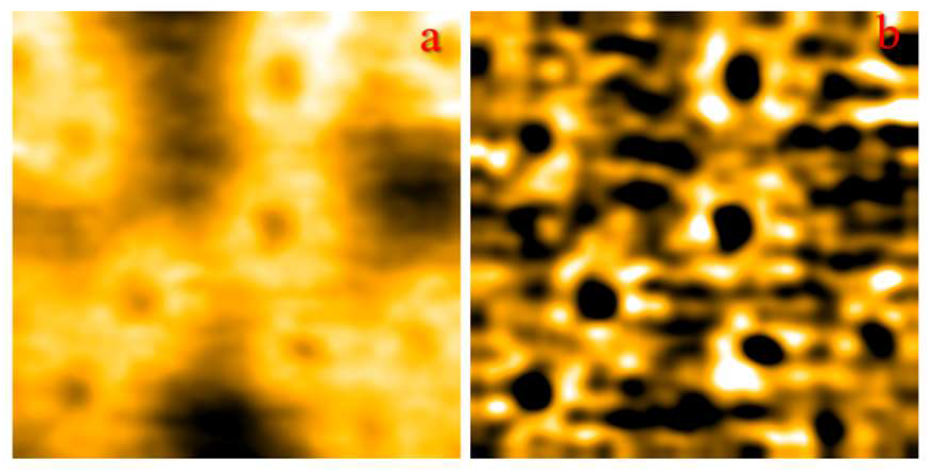
Example of filtering HS-AFM images using AFMnano3DR. (a) Original HS-AFM image; **(b)** Result after applying Gaussian and Laplacian filters to enhance αHL pore visibility.

**Fig. 8.**
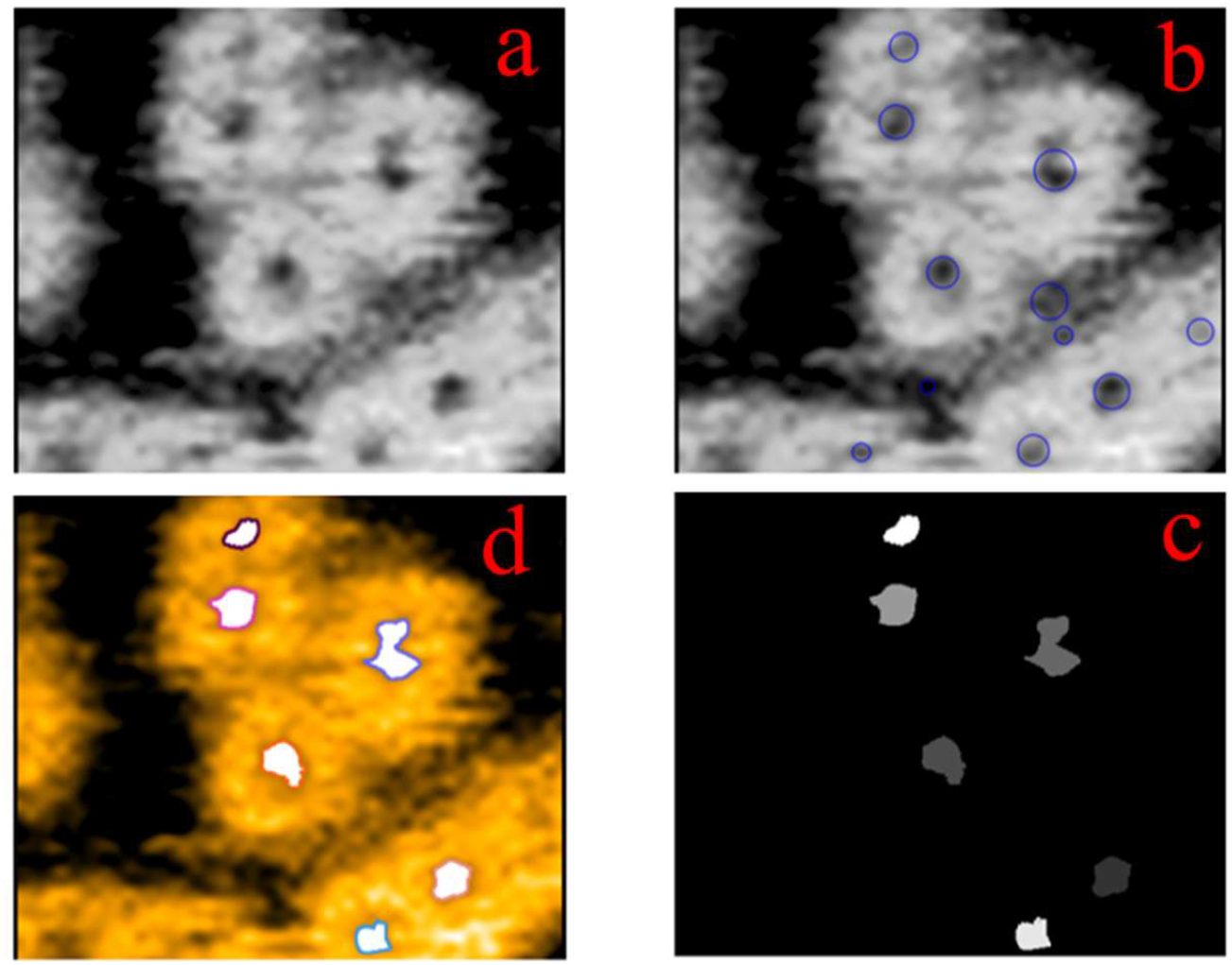
Illustration of instance boundary detection of internal pores in the pore stage of αHL using AFMnanoSALQ. **(a)** Input HS-AFM image; **(b)** Kernel detection identifies potential regions; **(c)** Kernel verification and curvature boundary extraction generate pores mask with unique IDs; **(d)** Pores are overlaid on the input image, each highlighted with a distinct color according to its mask ID.

**Fig. 9.**
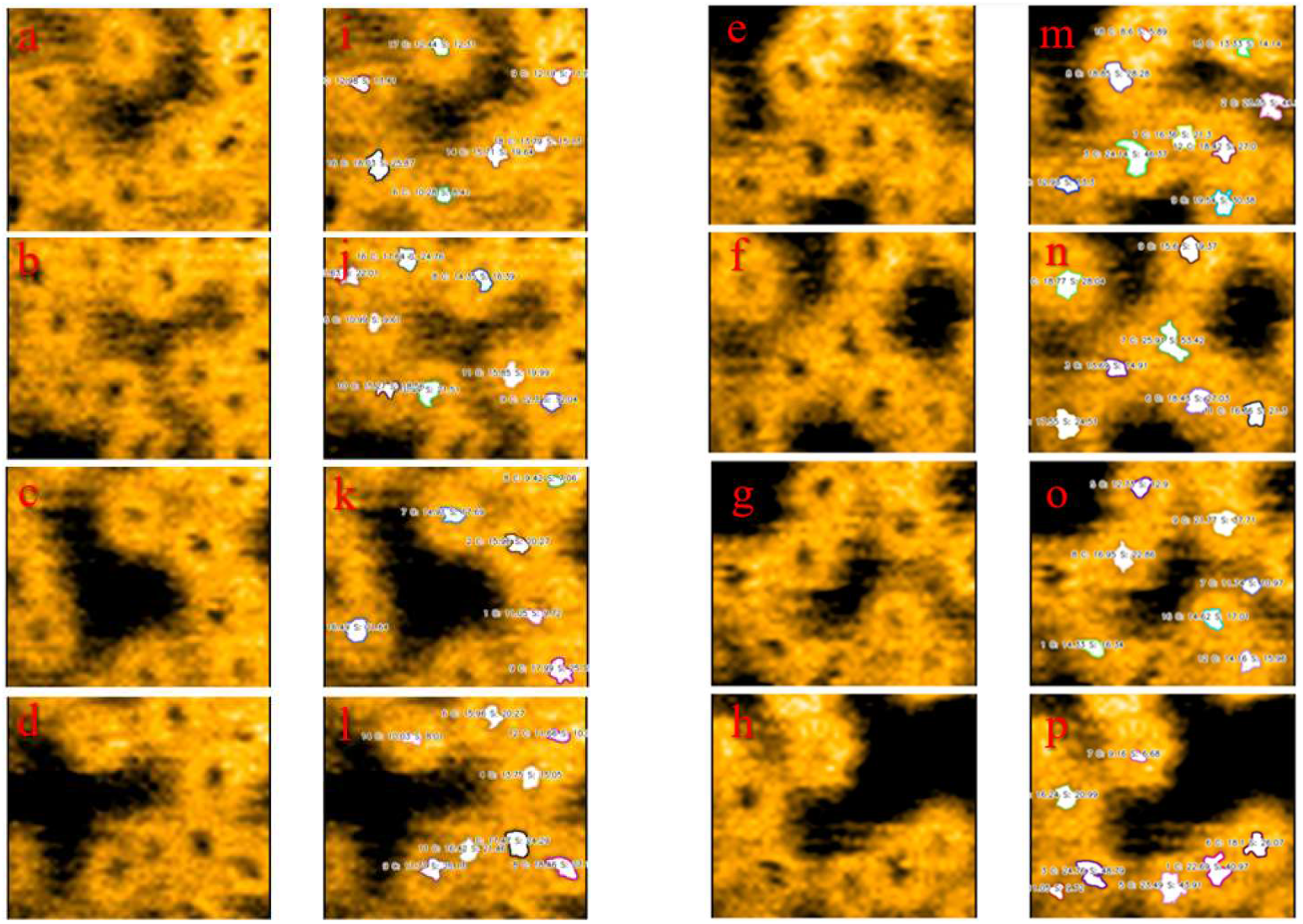
Illustration of instance boundary detection and quantitative analysis of αHL pores in HS-AFM images using AFMnanoSALQ. **(a)–(h)** Input HS-AFM images; **(i)–(p)** Corresponding results with detected pores labeled by instance. Quantitative measurements such as perimeter and area are overlaid for intuitive interpretation.

**Fig. 10.**
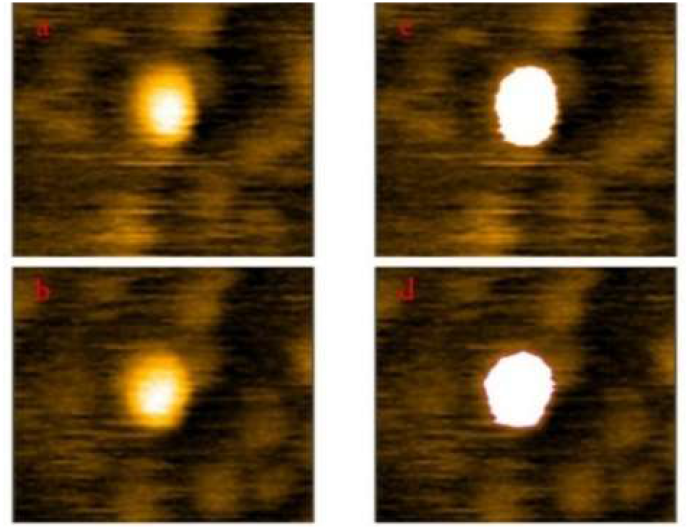
Illustration of instance boundary detection of αHL pre-pores in HS-AFM images using AFMnanoSALQ. **(a), (b)** Input HS-AFM images; **(c), (d)** Corresponding results with detected pre-pores labeled by instance.

**Fig. 11.**
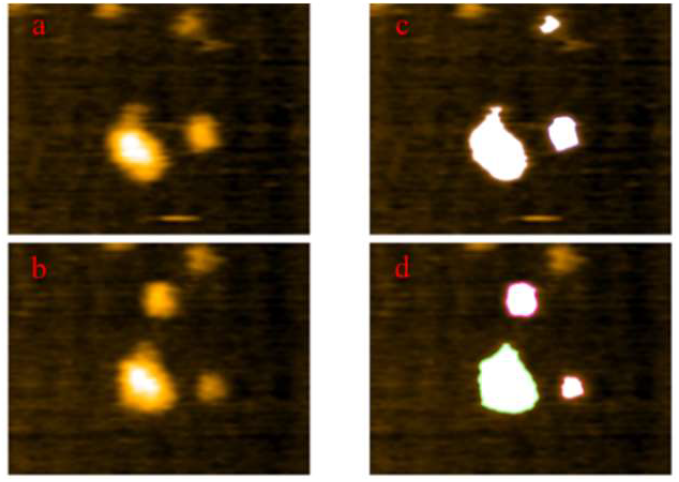
Illustration of instance boundary detection of oligomers at αHL intermediate stage in HS-AFM images using AFMnanoSALQ. **(a), (b)** Input HS-AFM images; **(c), (d)** Corresponding results with detected oligomers labeled by instance.

AFMnano3DR serves as a practical utility for tasks that require detailed inspection and manipulation of 3D topographic surfaces derived from AFM data, such as manual labeling, morphological analysis, or validation of segmentation results.

## Appendix 2. Example Outputs of AFMnanoSALQ

## Footnotes

1 RCSB PDB (https://www.rcsb.org/), PDBJ (https://pdbj.org/)

